# Engineering Biohybrid Mycelium Fibers through Hierarchical Structuring and Biomineralization

**DOI:** 10.64898/2025.12.17.694948

**Authors:** Olivia Pear, Kelly Balch, Jenna Stanislaw, R. Kōnane Bay

**Affiliations:** Materials Science and Engineering Program, University of Colorado Boulder, Boulder, CO 80303, USA; Department of Chemical and Biological Engineering, University of Colorado Boulder, Boulder, CO 80303, USA; Interdisciplinary Quantitative Biology Program, BioFrontiers Institute, University of Colorado Boulder, Boulder, CO 80303, USA

## Abstract

Driven by the persistence of microplastics and an overdependence on non-renewable sources, mycelium materials have emerged as alternatives to traditional materials, owing to their sustainable production and versatility. Engineered living materials, composite material systems incorporating biological components to enable function, possess desirable regenerative properties but have yet to be fully applied due to their lack of strength versus synthetic or natural materials. We leverage two strategies used by nature to generate mechanical strength: structural hierarchy and biomineralization. We extrude a bioink composed of alginate and a modified strain of the fungus *Aspergillus niger* capable of silica mineralization, and modulate the morphology by the growth and processing conditions. Mineralization results in significantly stronger and stiffer dried fibers. We demonstrate their potential as textile materials through twisting and braiding to significantly increase the fracture strain. Our results show that mycelium morphology and mechanics can be tuned through mineralization, growth conditions, and processing.

## Introduction

Textile materials are a significant source of primary microplastics, with synthetic clothing making up 35% of the microplastics released to oceans.^1^ It is estimated that 124 to 308 mg per kg, or 640,000 - 1,500,000 microfibers, are produced from one laundry cycle of synthetic textiles.^2^ According to the U.S. Environmental Protection Agency (EPA), the recycling rate for textiles was 14.7 % in 2018, presenting motivation to produce textile materials with reduced environmental impact and improved recyclability.^3^ Materials generated from fungi have the potential to displace incumbent textiles as they can be grown rather than extracted from finite resources.^4^ Despite a recent acceleration in startup creation and patent filing, materials generated from mycelia have been utilized by Indigenous people of North America for at least a century.^5^ Mycelia materials are utilized in industry for packaging, insulation, and sustainable leather.^6^ Research efforts have been primarily focused on composite mycomaterials, which derive their mechanical strength from the lignocellulosic biomass feedstock substrate that the fungus consumes.^7^ However, current mycelium materials lack strength compared to traditional polymer materials, necessitating exploration of mycelium in an aligned fiber morphology, which has, to the best of our knowledge, not been previously studied.^8^

Use of a microorganism for a textile material allows for inherent living properties such as growth and regeneration. The engineered living materials (ELM) field aims to design materials with specific functionality by capitalizing on the biological component.^9,10^ The use of living mycelia materials was first addressed by Gantenbein et al. by which they constructed mycelia robotic skin by three-dimensionally (3D) printing hydrogels loaded with the fungus, *Ganoderma lucidum*.^11^ Additionally, the self-healing functionality and recovery of mechanical properties after healing was addressed by Elsacker et al.^12^ Particularly due to their livingness and high water content, similar to synthetic hydrogels, these materials lack strength in the hydrated state. Though it is well known that many hydrogels increase strength upon drying, the design of ELMs is impacted by the loss in cell viability with dehydration. Mycelia can be dehydrated and remain living as metabolically dormant spores, providing a direct path for strengthening based on hydration state.^12,13^

One mechanism for generating strength and toughness in nature involves the creation of high impact-resistant materials through the formation of inorganic-organic composite structures by biomineralization. The interfacial binding of macromolecules and biominerals is critical in providing control over resultant structures and strength, exemplified by silicatein-α, an enzyme which forms silica in deep-sea sponge skeletons, and Pif, a key protein in calcium carbonate production for nacre formation.^14,15^ Intricate structural hierarchy is realized in the inorganic-organic twisted fiber composite of bone, the concentric layered glass and protein layers that comprise glass sponge spicules, and the cellular structure of wood. The structure and composition give rise to the toughness, flexibility, and strength of these respective materials.^16–19^ Sponge spicules, for example, are concentric lamellar structures composed of silica nanoparticles deposited around a central protein filament. Organic interlayers alternate with silica layers, serving to prevent crack propagation. Researchers are beginning to exploit nature’s approach to design stronger and tougher ELMs. For example, ELM nacre, designed as a bacterial planar composite, demonstrates compressive toughness within the range of natural nacre.^20^ However, we have yet to achieve the complex, hierarchical biomineralized structures observed in nature. Beyond planar materials, twist is observed in biological materials like collagen fibrils.^21,22^ This has been explored in composite living fibers, in which textile methods for processing staple fibers into yarn were used to replicate collagen twist.^23^

To develop hierarchical fibers, we leverage the filamentous fungal species *Aspergillus niger*. *A. niger* is responsible for at-scale production of citric acid and enzymes such as glucoamylases and phytases.^24^ The growth of a filamentous fungus occurs through apical extension of hyphae, which are tubular-like structures with a diameter between 2-10 µm. Hyphae form extended networks (i.e., mycelia) through branching and recombining with neighboring hyphae.^25^ The growth mechanism of hyphae within a mycelium network can be shifted between phalanx and guerilla foraging, to balance exploitation or exploration in nutrient-dense and nutrient-deficient areas, respectively, by manipulating nutrient availability.^26^ The cell wall of the hyphal tip contains primarily polysaccharides such as alpha- and beta-glucans (50-60%) and chitin (10-20%).^27^ The polysaccharide components of the fungal cell wall, namely beta-glucans and chitin, confer strength to the mycelium.^28–31^ The presence of chitin impacts the cell wall stiffness, contributing to an elastic modulus ranging from 10-100 MPa.^32^ This range in moduli is due to apical growth, wherein the cell wall rapidly accumulates the expansion and generation of more cell wall material, contributing to a variation in thickness from 50-500 nm.^32^ All combined, we hypothesize that hierarchical alignment of mycelia is possible across multiple length scales by nanoscale extracellular enzymatic silica production, micron-scale hyphae filament growth, and macroscale structure control by application of twist and braid.

In this work, a mycelium bioink is extruded to produce mm-scale fiber structures with micron-scale features. The fungus, *A. niger*, is engineered to produce nanoscale mineral deposits through extracellular enzyme activity. The mycelia fiber can be tested as a single fiber and may additionally be twisted or braided into a more complex macroscale structure. Mineralized mycelia demonstrate increased strength compared to non-mineralized mycelia. Further, twisted and braided fibers demonstrate an increase in fracture strain compared to single fibers. Overall, these results provide insights into how biomineralization and hyphal alignment impact the mechanical properties of mycelia ELMs.

## Results and Discussion

### Extrusion of living fungal fibers

To investigate the tensile properties of an aligned mycelia network, a fiber morphology was chosen to confine the fungus in a crosslinked structure, allowing for radial hyphal growth. Fibers, approximately 1 mm in diameter, were created by the extrusion of a bioink composed of sodium alginate and *A. niger* fungal cells into a calcium chloride bath (Figure 1). Alginate, a shear-thinning anionic polymer, reversibly crosslinks upon interaction with divalent or trivalent ions, which are supplemented into fungal nutrient media by calcium chloride. Thus, alginate additionally serves as a sacrificial scaffold.

**Figure 1.**
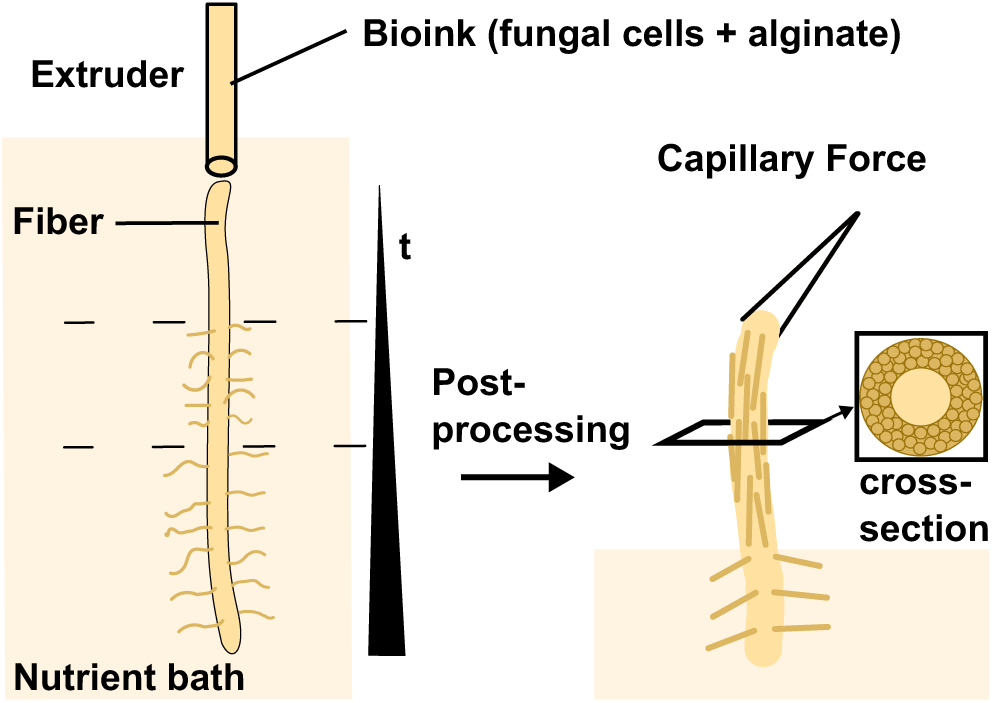
Fungal bioink extrusion and alignment. Schematic demonstrating the extrusion of fungal bioink into a nutrient bath, radial growth of fungus by hyphae, and subsequent collapse of hyphae to the fiber core upon removal from bath.

The ink instantaneously crosslinks after extrusion through the nozzle in the calcium chloride bath, combining ease of extrudability with fixing of printed structure through gelation. The fungal cells include both 5 µm diameter spores and approximately 0.5 mm diameter mycelia pellets. During extrusion of the bioink, we observe an increase in toughness (*P* < 0.05) of pure alginate fibers extruded through a smaller diameter 20 gauge needle (internal diameter: 0.6604 mm) compared to a larger diameter 18 gauge needle (internal diameter: 0.9652 mm). (Figure S1). The increase in toughness with decrease in needle diameter is attributed to the alginate polymer becoming aligned due to shear stress experienced in the needle. An 18 gauge needle is used to extrude ink containing mycelia pellets. Additionally, using a combination of fungal cells at different growth stages is beneficial to the network formed as the mycelia pellets are more mature and exhibit rapid growth, while the spores minimize void space.

### Effect of hyphal growth on strength

Suspension of the fibers into the liquid nutrient bath supports radial fungal growth by hyphal filaments. We first determined the optimal growth conditions required for hyphal extension. Multiple fibers were suspended in a nutrient bath at 30 °C, and hyphal length and biomass were measured every 24 hours. At 30 °C, we observed a plateau in the hyphal extension of encapsulated fungal cells in gelled alginate fibers after 72 hours of growth; therefore, 72 hours was chosen as the control growth time (Figure S2). The growth conditions influence the rate of hyphal extension, the length of hyphae, and the density of hyphae. The introduction of shaking conditions during growth, by incubating a single fiber on a stir plate to aerate the medium, resulted in denser hyphae, whereas stationary growth of a single fiber resulted in sparser hyphae (Figure 2A). We associate the difference in growth morphology to the availability of nutrients. High availability of nutrients due to continual mixing results in phalanx-type foraging of mycelia, a more uniform growth of fungus that results in denser hyphae. Stationary growth leads to nutrients at the cell periphery becoming depleted more readily, and mycelia utilize more guerilla-type foraging, evidenced by sparser, branched hyphae. At room temperature, we observe a lack of significant fiber hyphal growth until 72 hours. By 168 hours, single fibers grown under static conditions have longer hyphae compared to shaking growth (Figure 2B).

**Figure 2.**
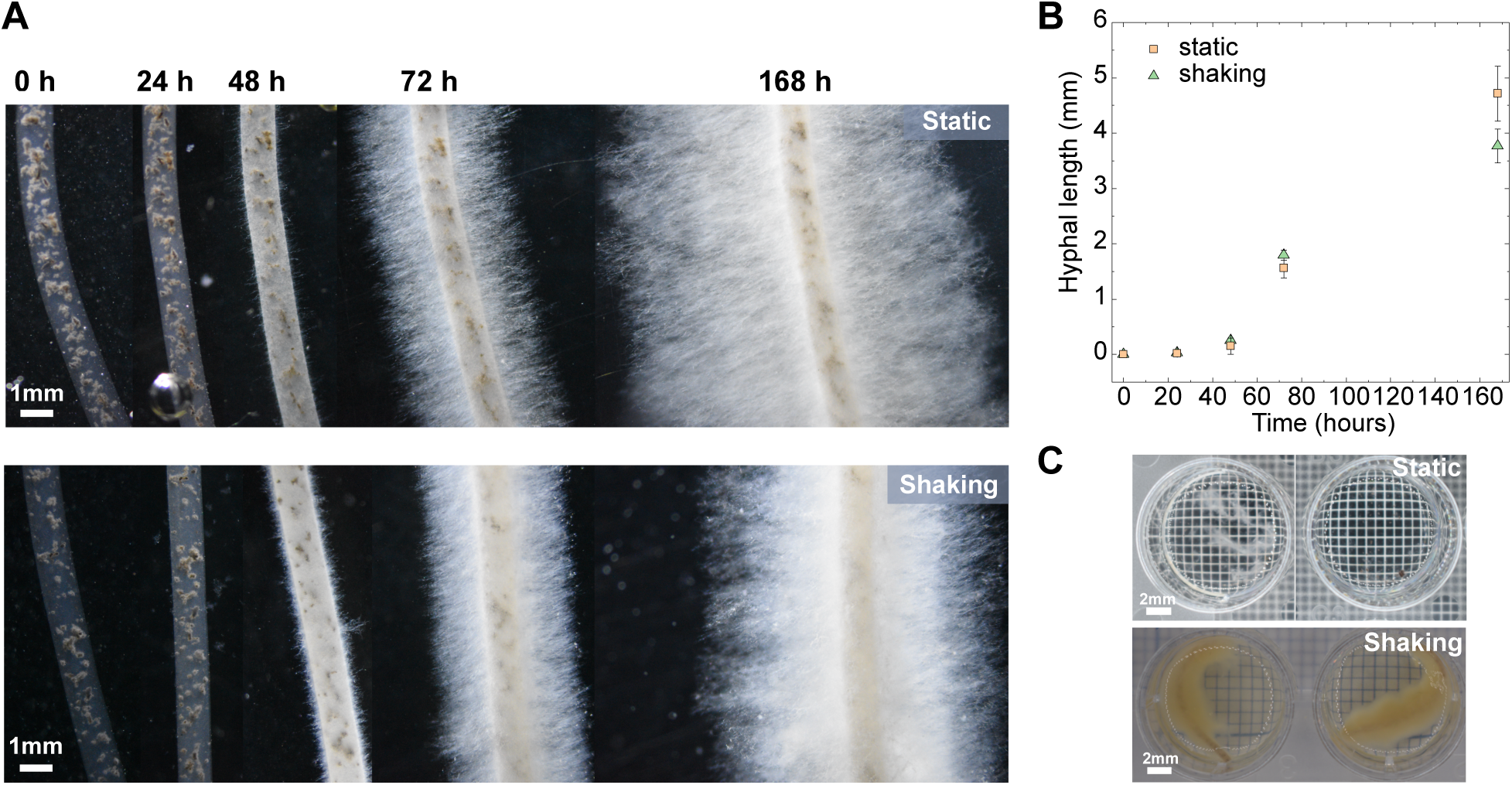
Fungal growth behavior and sacrificial support degradation. A) Zoom lens photographs of hyphal radial growth at room temperature, under static (top row) or shaking (bottom row) incubation. Images are representative sections of the fiber at each time point. B) Hyphal length measured over time for static and shaking growth conditions. The data represent the mean values and the standard deviation of five different samples. C) Mycelia fiber, from growth vessel with multiple fibers, after alginate de-crosslinking for static and shaking growth with two different samples shown for each condition.

In order to investigate whether the mycelia formed a network internal to the fiber core, we dissolve the alginate structure using a chelating agent, ethylenediaminetetraacetic acid (EDTA). Notably, the removal of the alginate network from the fiber by depolymerization demonstrates a lack of internal mycelia structure in static growth samples, while the fiber structure of samples grown under shaking conditions remains intact (Figure 2C). We note this distinction in network structure is observed only for multiple fibers grown in the same media. With single fibers grown separately, we observe intact fibers after incubating with EDTA beginning at 48 hours for both static and shaking growth, suggesting that internal mycelia network formation may depend on nutrient competition. We hypothesize that the shaking condition results in increased growth-based crosslinking within the fiber core.

The growth conditions provide further control of alignment across multiple length scales. The variation in hyphal morphology due to growth conditions also impacts fiber structure upon removal from the nutrient bath, assessed by scanning electron microscopy (SEM, Figure 3). After stationary growth, collapsed hyphae are largely aligned, and capillary alignment of hyphae can be observed (Figure 3, middle). Sparse hyphae readily collapse to the core fiber. However, dense hyphae, such as those formed during shaking growth, retain their extended structure and avoid full collapse to the fiber core. This is evidenced by the broad fiber thickness and random hyphae network (Figure 3, right).

**Figure 3.**
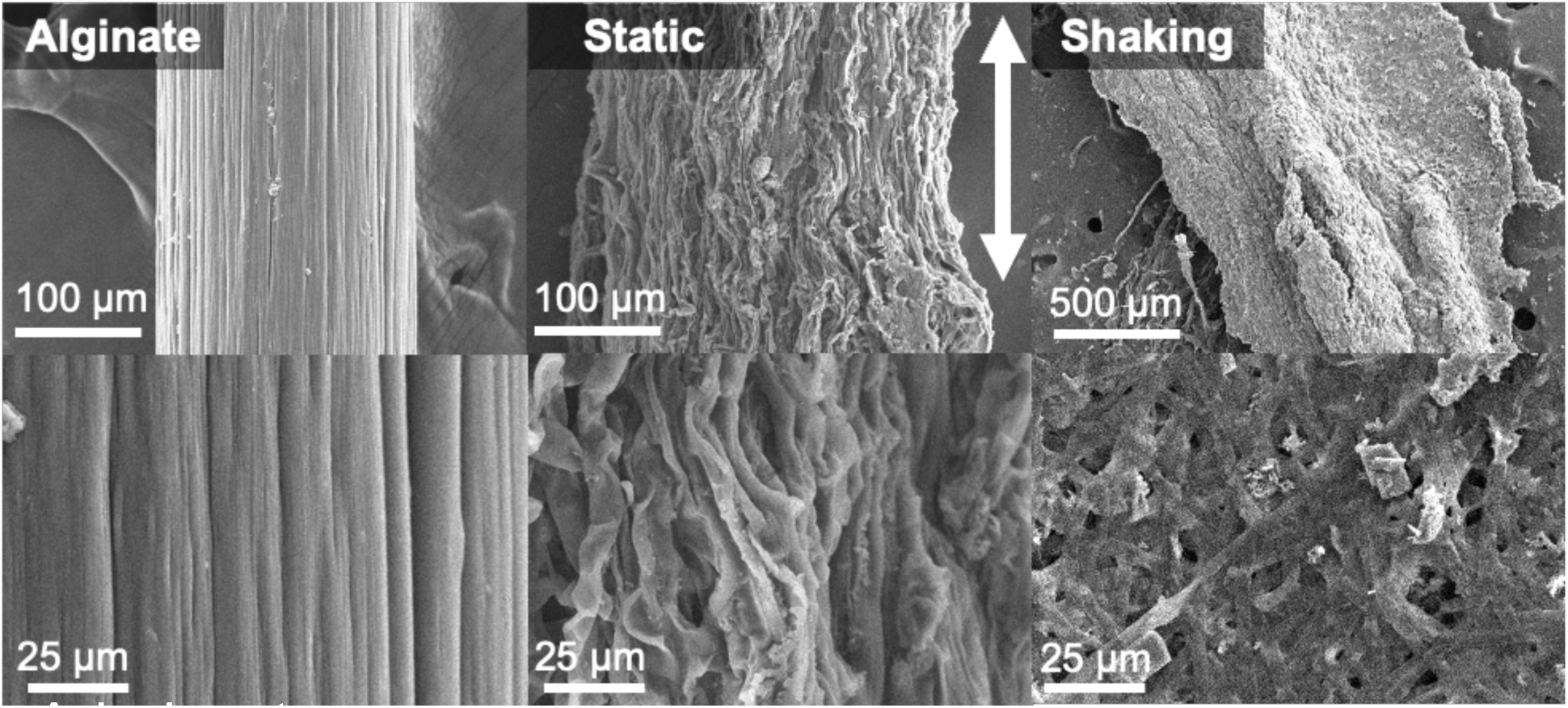
Alignment of mycelium under varied growth conditions. Scanning electron microscopy images of pure extruded alginate (left column), extruded mycelia fiber incubated statically (middle column), and extruded mycelia fiber incubated under shaking conditions (right column).

We hypothesize that the observed hyphal alignment along the axis of the fiber core contributes to the overall strength of the fiber, as shown in studies of composites with varied length and orientation.^33^ To evaluate the hyphal alignment contribution to the strength of the fiber, we perform tensile test measurements on hydrated samples incubated for varied amounts of time. We observe that the elastic modulus, *E,* of hydrated fibers grown statically or with shaking increases significantly to 420 kPa by 72 hours of growth, which we associate with increased hyphal alignment due to growth and subsequent capillary force collapse (Figure 4). We observe a subsequent decrease in modulus after 120 h of growth to values comparable to those at 0 hours of growth. This observed loss in crosslinked alginate structure after 72 hours of growth is attributed to the reduction in pH of the nutrient bath. The production of organic acids by the fungus, such as oxalic acid and citric acid, is known to contribute to the reduction in growth medium pH.^34^ Our prior work demonstrates a reduction in pH of the culture medium over the course of growth.^35^ We hypothesize that the acidification of the media leads to weakening of the fiber in the hydrated state, as the fiber properties are alginate-dominated.^36^

**Figure 4.**
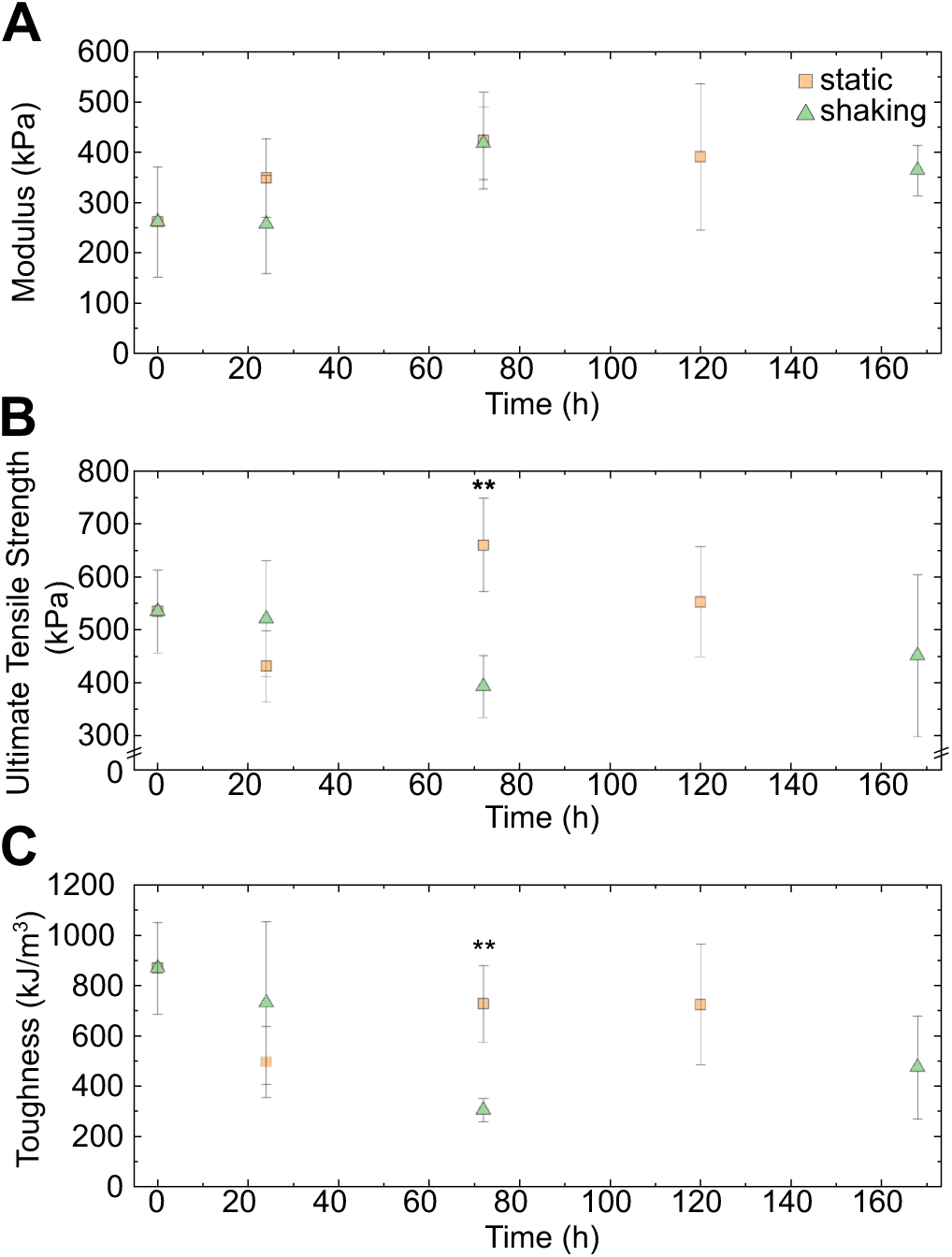
Tensile mechanical properties of hydrated fibers. A-C) Modulus, ultimate tensile strength, and toughness of hydrated fiber samples incubated with static or shaking growth. The data represent the mean values and the standard deviation of five different samples. ** *P* < 0.01, one-way ANOVA followed by Tukey’s *post hoc* test comparing static and shaking growth. Statistical significance of growth times is determined by one-way ANOVA with Tukey’s *post hoc* test and p-values and data are reported in Table S1.

Similarly, fibers grown under shaking conditions exhibit an increase in modulus between 24 h and 72 h of growth and subsequent decrease after 168 h of growth. Static growth shows an increase in ultimate tensile strength between 24 h and 72 h, returning to 0 h values by 120 h growth. For the hydrated fibers grown under shaking conditions, the ultimate strength does not significantly vary across growth from 0 h to 168 h. At 72 h, we observe higher ultimate strength of static growth compared to shaking growth. The statically grown fibers exhibit a significant drop in toughness at 24 h, which plateaus for the remaining time points tested (up to 120 h) and returns to 0 h values by 72 h growth (Figure 4). Additionally, toughness decreases from 0 h to 168 h of growth for fibers grown under shaking conditions, with substantial drop in toughness at 72 h. At 72 h, the decrease in toughness is also observed between the growth conditions, with static growth exhibiting higher toughness compared to shaking growth. Lower strength and toughness of the fibers in the hydrated state is attributed to the weakening of calcium crosslinking by the lower pH environment as well as fungal growth physically disrupting the alginate structure. The elastic modulus of fibers in the hydrated state is impacted by growth and hyphal length, increasing up to 72 h of growth and then plateauing. The modulus in the kPa regime is reflective of the gelled alginate biopolymer, with the presence of the fungus resulting in increased strength relative to pure 3 wt. % alginate (Figure S3).^37^

### Biomass and composition

To further understand the impact of processing conditions on growth, we measure dry fiber mass. Single fibers are grown in tissue culture flasks for varying amounts of time. Fibers are dissected into 1 cm fragments and mass recorded after drying overnight in an oven at 60 °C. We observe an increase in dry fiber mass after 72 h growth as well as a distinction in dry fiber mass between static and shaking growth beginning at 72 h, and most prominently, at 168 h (Figure 5).

**Figure 5.**
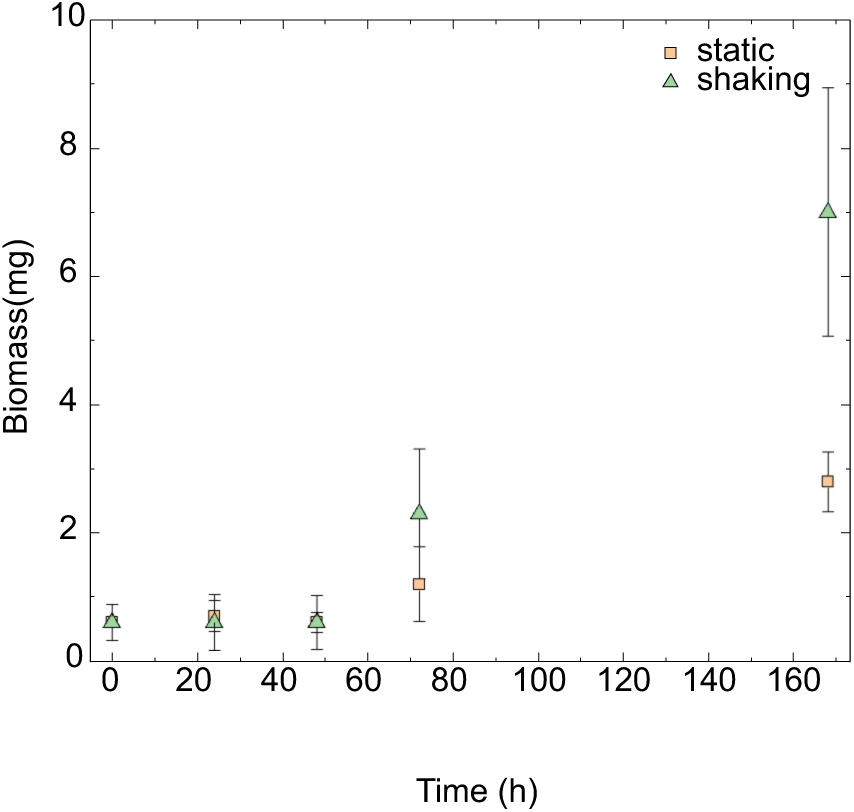
Growth of single mycelia fibers in complete media. Amount of biomass generated by the fungus over time by static (orange, square) or shaking (green, triangle) growth conditions. The data represent the mean values and the standard deviation of five different samples.

In addition to biomass measurements, we use Attenuated Total Reflectance-Fourier Transform Infrared Spectroscopy (ATR-FTIR) to detect the presence of functional groups within fungal samples prepared with static and shaking growth conditions and to establish the relationship between hyphal morphology and biochemical structure. In statically grown samples, FTIR spectra show defined medium peaks at 1400 cm^-1^ and 1600 cm^-1^ and a strong peak at 1050 cm^-1^, corresponding to amide bands and a distinct C-O peak identified in chitin, respectively (Figure S4).

In samples grown under shaking conditions, there are weaker and less defined peaks at 1400 cm^-1^ and 1600 cm^-1^, corresponding to a reduction in amide bands. Additionally, the presence of new peaks at 1370 cm^-1^ and 1200 cm^-1^ suggests an increase in polysaccharide production in the fungus grown with shaking conditions. Finally, we observe an increase in weaker peaks at 2800-2900 cm^-1^, suggesting an increased presence of lipids in the fungus with shaking growth. Our results are in agreement with prior studies, which found that agitation enhanced biomass growth and exopolysaccharide (EPS) production of submerged fungal species *Paecilomyces sinclairii* and *Paecilomyces tenuipes*.^38,39^ Additionally, low shear rate mixing resulted in mycelium pellets with a fluffy perimeter of hyphae.

### Fiber mineralization

Our approach to address the tensile strength limitations of mycelia materials includes the investigation of the effect of nanoscale enzymatic mineralization on macro-scale mechanical strength. In nature, biomineral deposits enhance the strength and toughness of many composites, such as bone, nacre, and sea sponges.^40,41,18^ Building on our prior work, we use a genetically engineered strain of *A. niger* fungus that expresses extracellular silicatein enzyme from the marine sponge *Latrunculia oparinae* to enable surface silica coating.^35^ To facilitate mineralization, we suspend the fibers in an aqueous solution containing 10 mM hydrolyzed tetraethyl orthosilicate (TEOS) at pH 7 for 24 hours after an initial growth period of 72 hours.

When studied in the hydrated state, 72 h statically-grown fibers exhibit a weakening effect when exposed to TEOS, reflected in lower ultimate tensile strength (UTS) (*P* < 0.001) and toughness (*P* < 0.001) (Figure S3). The modulus of fibers treated with TEOS is not significantly different from untreated fibers. Pure alginate hydrated fibers incubated with TEOS also did not exhibit any difference from pure alginate at neutral pH. We observe a decrease in fiber toughness (*P* < 0.05) and modulus (*P* < 0.05) of pure alginate fibers incubated with TEOS at pH 2.4 relative to pH 7.6 (Figure S3). The decrease in toughness of fungus-containing alginate fibers over the course of growth is, thus, attributed to the reduction in pH of the nutrient bath.

In the dried state, however, changes in the properties of the biomineralized mycelium network are observed. Biomineralization of dried extruded fungal cell fibers increases strength (*P* < 0.01) and modulus (*P* < 0.01) relative to the non-mineralized control, which we associate to the silica deposition from the enzymatic activity of silicatein-𝛼 in catalyzing the mineralization (Figure 6). Such mineralized, dried single fibers demonstrate a modulus on the order of 1 GPa and ultimate strength of 40 MPa, a 3 - 4-fold increase compared to non-mineralized single fibers. Our previous work demonstrated the presence and quantification of silica from enzymatic mineralization of *A. niger* films using scanning electron microscopy with coupled energy dispersive X-ray spectroscopy, thermogravimetric analysis, and uniaxial tensile testing.^35^ The degree and type of mineralization can be modulated by using other metals as substrates, including titanium and gallium, as silicatein has been shown to have non-specific catalytic activity.^42,43^ The modularity of our system could be tested in the future by swapping substrates to generate conductive fibers from mycelium. The modulus and strength for mineralized mycelium films are much weaker relative to *A. niger* fibers, at 2.5 MPa and 325 kPa, respectively, which we attribute to the lack of hyphal alignment in the mycelium network.^35^

**Figure 6.**
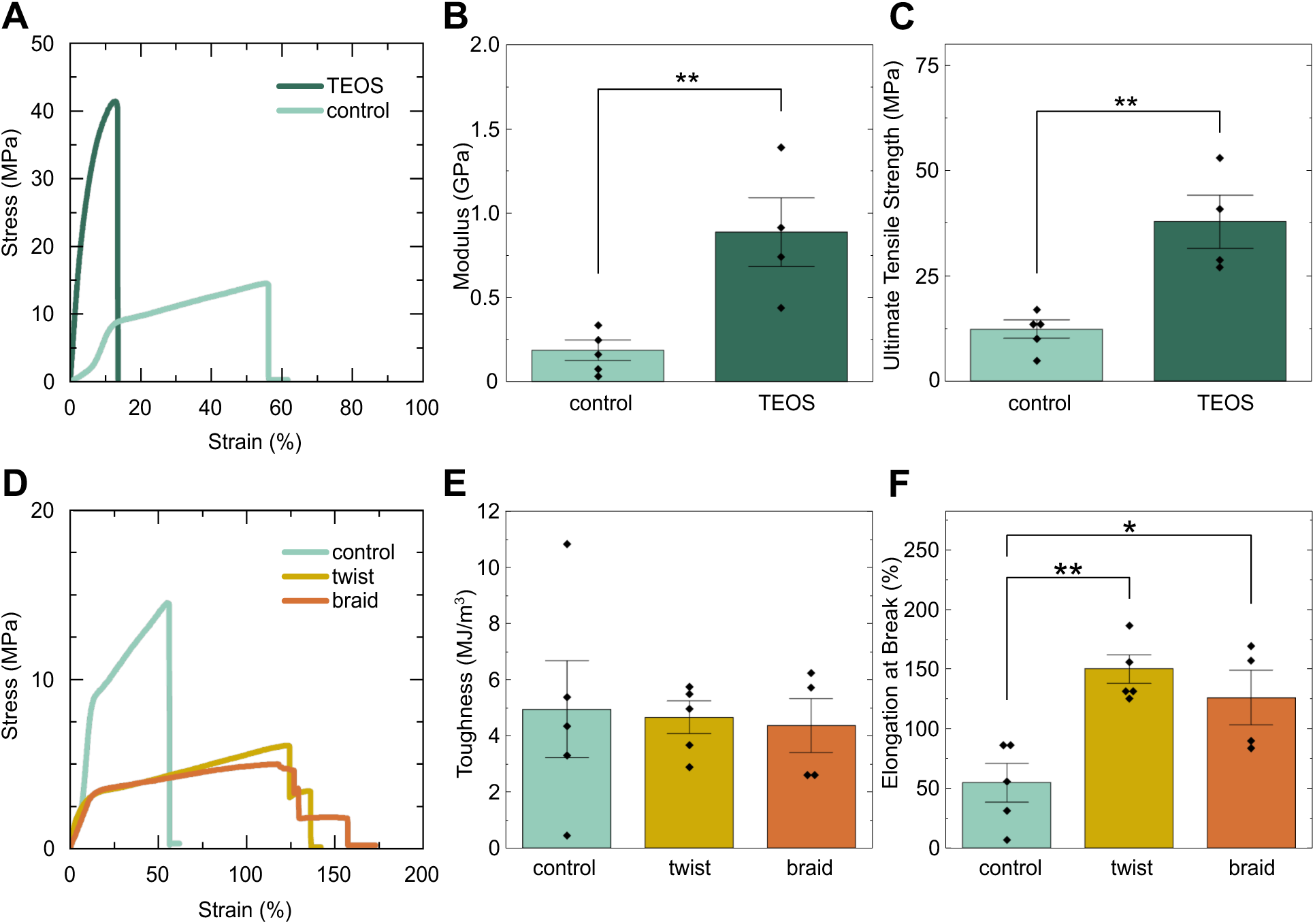
Tensile mechanical properties of dried fibers. A) Representative stress-strain curves of untreated and TEOS treated dried mycelia fibers obtained from uniaxial tensile testing. B,C) Modulus and ultimate tensile strength of untreated and TEOS treated dried mycelia fibers. The bars represent the mean values and standard error of the mean for the individual data points shown. D) Representative stress-strain curves of twisted and braided dried mycelia fibers compared to single dried fiber. E,F) Toughness and elongation at break of twisted and braided mycelia fibers. The bars represent the mean values and standard error of the mean for the individual data points shown. * *P* < 0 .05, ** *P* < 0.01, one-way ANOVA followed by Tukey’s *post hoc* test.

Given the lack of recyclability and generation of microplastics with current textile production and usage, it is of interest to compare our mycelia fibers to the mechanical properties of a synthetic textile fiber, elastane. Uniaxial tensile testing of a 55 denier elastane fiber, with an average diameter of 60 μm, results in a modulus of 0.015 GPa and tensile strength of 62 MPa (n=3). 800 denier elastane, with an average diameter of 300 μm, results in a modulus of 6 MPa and tensile strength 20 MPa (n=1) (Figure S5). The mycelia fibers exceed the modulus of elastane, and mineralized mycelia fibers approach the tensile strength of 55 denier (den) elastane.

### Fiber twist

In nature, twisted architectures serve to improve mechanical properties and biological function.^44^ Natural twist, like that observed in wood and bone, and applied twist, achieved by spinning fibers into yarns increase strength by resisting torsional force. In yarns, torque is generated by twisting fibers together, leading to individual fibers becoming more compact. To impart additional strength in the extruded fibers, we aim to direct the macro-scale alignment using techniques such as manual twisting and braiding.

Braided and twisted samples are generated while hydrated and then dried before testing. We observe a significant increase in fracture strain, or elongation at break, for twisted (*P* < 0.01) and braided (*P* < 0.05) samples compared to single fibers with fracture strain increasing from 50 % to over 100 % in both twisted and braided samples (Figure 6F). The shape of the representative stress-strain curves suggests a toughening mechanism indicated by multi-stage fracture (Figure 6). Twisted and braided fibers show significantly lower ultimate strength but comparable toughness and modulus to single fibers. The overall mechanical properties result from a balance of fiber compactness and friction, which contributes to increasing strength, and the obliquity of component fibers to the yarn axis, which contributes to decreasing strength.^44^ We hypothesize that the lower strength observed arises from weak twisting, allowing the fibers to straighten more under tension, with limited inter-fiber cohesion that would cause an increase in strength. Future studies could investigate the effect of twist angle, twist density and the bundling of multiple fibers to improve inter-fiber cohesion and strength.

## Conclusion

With the increase in interest to create functional living materials that approach the mechanical properties of conventional materials, extrusion-printed fungal cells have emerged as a promising platform. Containing the cells in a crosslinked alginate network structurally supports the fungus while enabling mycelia network development by radial growth towards a nutrient source. By modulating the growth environment and controlling mineralization, we generate structures with enhanced strength and stiffness, as well as diverse morphologies across multiple length scales. We anticipate that the increased strength and processibility, coupled with the living nature of the mycelia, enables the design of mechanically-robust ELMs that self-assemble, adapt, and regenerate.

## Methods

### Fungal Strain

The *Aspergillus niger* strain with surface-expressed silicatein (SilA) used in this work was constructed and provided by the Khakhar lab at Colorado State University and has been previously described.^35^ Briefly, Golden Gate Assembly was used to construct plasmids encoding expression cassettes for SilA using the Modular Synthetic Biology Toolkit for Filamentous Fungi.^45^ A fungal entry vector containing homology regions to the pyrG locus was constructed to enable homology directed repair mediated knock-in. The entry vector and expression cassette were assembled together, sequence verified, and transformed into *A. niger* using protoplast mediated transformation.

### Fungal Growth Conditions

Cultures were grown in Complete Media for *Aspergillus* as described.^46^ Complete media was supplemented with the described Vitamin Supplement. For incubation in TEOS, hydrolyzed TEOS was prepared as described.^47^ Hydrolyzed TEOS was added to water to a final concentration of 10mM and pH adjusted to 7.

Spores are generated from frozen glycerol stock on complete media agar plates, and grown for 5-7 days, until sporulation occurs on mycelia surface. To culture mycelia pellets, a roughly scraped surface of a sporulated agar plate is added to a 250 mL flask containing 100 mL complete media on an orbital shaker at 150 rpm and 30 °C for 20 hours. The nutrient bath for printing consists of complete growth media supplemented with 2 wt.% calcium chloride.

### Bioink Preparation

Mycelium inks for extrusion were prepared by first dispersing sodium alginate (5.5 wt. %) in sterile water and mixing on a stir plate until dissolved. Spores are harvested from an agar plate by gently pipetting water onto the surface until the concentration of the collected stock is above 10^8^ spores mL^-1^. To an ink volume of 15 mL, 9 mL of 5.5 wt. % sodium alginate, 10^8^ spores mL^-1^, and 4.5 mL of mycelia pellets are added, and adjusted to the final volume with sterile water. The ink is briefly mixed with a spatula before use.

### Bioink Extrusion

Fibers are extruded using Lulzbot Mini2 extrusion printer modified to Replistruder 4 according to established protocol, operated inside a biosafety cabinet.^48^ The ink is loaded into a 3 mL syringe and extruded with a 18 G needle 1 inch in length. The bioink is printed directly into a nutrient media bath supplemented with 2 wt.% calcium chloride to ionically crosslink the sodium alginate upon contact with the fluid. Fibers are vertically suspended from screws using binder clips to allow for radial hyphae growth for up to 168 hours. Pictures of the growth were taken using a Nikon D3500 camera with ThorLabs 12.5X zoom lens.

During growth, fibers are incubated under stationary conditions or mixed on a stir plate at 150 rpm at room temperature unless otherwise stated. Biomineralized samples are exposed to 10 mM hydrolyzed TEOS in water at pH 7, unless otherwise stated. Three strands or two strands of fibers in the hydrated state are intertwined after growth to generate braided or twisted samples. Samples are dehydrated by conditioning overnight in a 55 % relative humidity chamber, using a saturated solution of magnesium nitrate.

### Mechanical testing

The mechanical properties were measured by uniaxial tension tests using a TA.XT*Plus*C Texture analyzer (5 kg load cell, Texture Technologies). Tests were conducted at a constant displacement rate of 0.1 mm sec^-1^. Samples were cut from original fiber length to approximately 10 mm and were mounted onto a paper frame with 5 mm gage length. The frame was mounted between the instrument grips and each side cut after alignment and tightening. Sample diameters were measured through image analysis using ImageJ/FIJI and used to determine cross-sectional area for calculating stress. Force-displacement data generated by the instrument were converted to stress-strain data using Matlab code. The elastic modulus was determined from the linear region of the stress-strain curve, between 1 - 5 % strain for hydrated fibers or 0.5 – 1 % strain for dried fibers. The ultimate tensile strength, UTS, was determined from the maximum stress of the specimen. The toughness was determined as the area under the stress-strain curve up to the UTS. Fracture strain was determined as the strain at maximum stress.

### Biomass Measurements

The biomass generated by printed mycelia fibers during growth was determined. Weighed samples were placed in 1.5 mL microcentrifuge tubes, where 100 µL of 0.1 M EDTA was added to the sample to de-crosslink the calcium alginate polymer network. Samples were incubated at room temperature for a minimum of 4 hours, centrifuged, supernatant removed, and subsequently washed with Milli-Q water and re-centrifuged. The pelleted sample was dried in an oven set at 60 °C overnight and weighed.

### FTIR

Samples were dried at room temperature for at least 48 hours in 24-well plates and ground to a fine powder before measuring. ATR-FTIR (Thermo Scientific Nicolet iS20 with a Ge Crystal ATR accessory) was performed using 64 scans at a resolution of 4 cm^-1^.

### SEM

Samples were removed from the tensile frame and dried at room temperature for at least 48 hours on an SEM stub prepared with carbon tape, in a petri dish. Before imaging, samples were sputter-coated (Cressington 108 Auto/SE) with 4 nm of platinum (Pt). Samples were imaged with a Hitachi TM-4000PlusE-2 with 15 kV accelerating voltage and 10 mm working distance. Elemental mapping was performed at 15 kV with an energy dispersive spectroscopy detector (Oxford Instruments).

## Acknowledgements

The authors thank Dylan Moss and Arjun Khakhar (CSU Department of Biology) for providing the fungal strains. The authors thank Natalie Fisher and Cécile Chazot (Department of Materials Science and Engineering, Northwestern University). The authors thank Matthew Fyfe for assistance with FTIR measurements carried out in the Living Materials Lab. SEM was performed at COSINC-CHR CU Boulder (RRID:SCR_018985). Funding: This work was funded by National Science Foundation (CBET 2516055), and CU Boulder startup funds. O.P. acknowledges support from Graduate Assistance in Areas of National Need (GAANN) fellowship award. J.S. acknowledges support from Interdisciplinary Quantitative Biology (IQ Biology) Program at the BioFrontiers Institute, University of Colorado, Boulder.

## Author Contributions

Conceptualization, O.P. and R.K.B.; methodology, O.P.; formal analysis, O.P. and R.K.B.; investigation, O.P., K.B., and J.S.; data curation, O.P.; writing – original draft, O.P. and R.K.B.; visualization, O.P.; writing – review & editing, O.P. and R.K.B.; software, O.P.; resources, R.K.B.; supervision, R.K.B.; project administration, R.K.B; funding acquisition, R.K.B.

## Resource Availability

### Lead Contact

Requests for further information and resources should be directed to and will be fulfilled by the lead contact, Kōnane Bay.

### Materials Availability

All materials generated in this study are available from the lead contact upon reasonable request.

### Data and Code Availability

Data provided as supplemental information and code is available on github.

