## Supplement information for "Engineering Biohybrid Mycelium Fibers through Hierarchical Structuring and Biomineralization"

Supplemental Figures

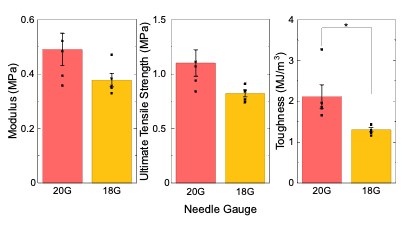

**Figure S1. Alginate fiber tensile properties.** Modulus, ultimate tensile strength, and toughness of 20 gauge (pink) and 18 gauge (yellow) extruded alginate fibers tested by uniaxial tensile testing. The bars represent the mean values and standard error of the mean for the individual data points shown. * *P* < 0.05 by Student’s t test.

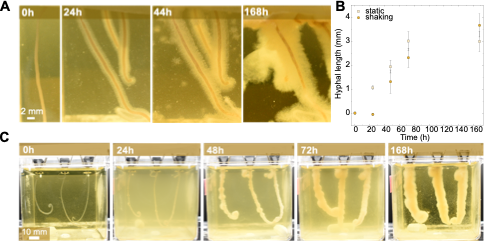

**Figure S2. Growth behavior of multiple mycelia fibers grown together.** A) Photographs of mycelia fibers grown under static growth conditions with increasing growth time. B) Hyphal length measured over time for static and shaking growth conditions. The data represent the mean values and the standard deviation of five different samples. C) Photographs of mycelia fibers grown under shaking conditions with increasing growth time.

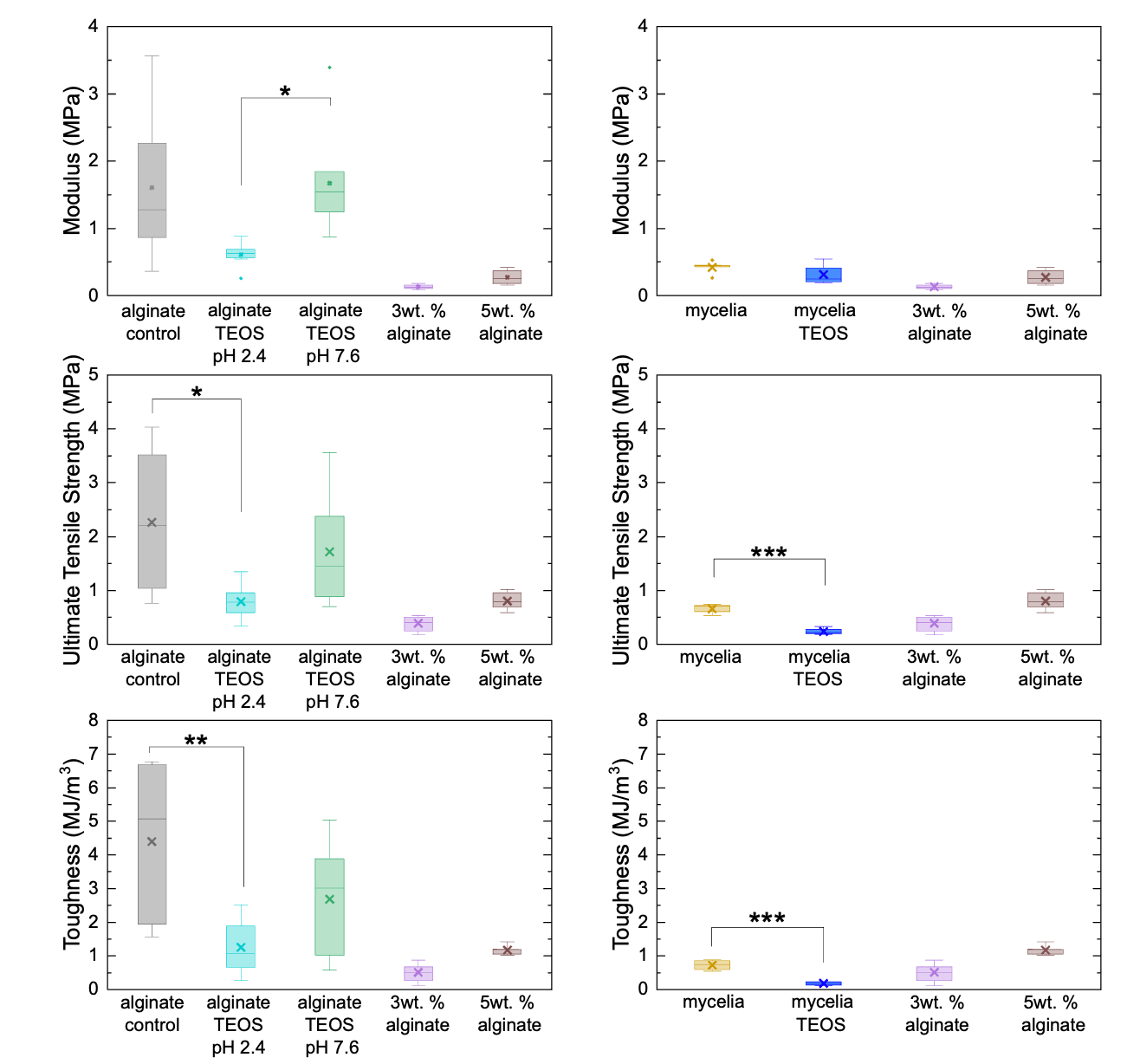

**Figure S3. Effect of incubation, substrate, and fungus on tensile properties relative to pure alginate fibers.** Modulus, ultimate tensile strength, and toughness of alginate fibers tested by uniaxial tensile testing. Alginate fibers incubated at 30 °C for 24 hours (n=7, grey), or incubated in TEOS at either pH 2.4 (n=8, cyan) or pH 7.6 (n=7, green). 3 wt. % alginate (n=7, purple) and 5 wt. % alginate (n=7, brown) serve as controls tested after fiber generation. Mechanical strength of mycelia fiber (n=5, yellow), mycelia with TEOS (n=4, blue), with the same controls 3 wt. % (n=7, purple) and 5 wt. % (n=7, brown). * *P* < 0.05, ** *P* < 0.01, *** *P* < 0.001, one-way ANOVA followed by Tukey’s *post hoc* test.

**Table S1. *p*-values of tensile properties of hydrated fibers.**

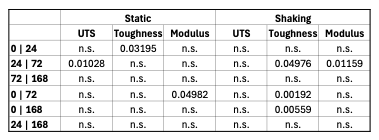

Statistical significance of growth times in static and shaking conditions determined by one-way ANOVA with Tukey’s *post hoc* test. n.s. not significant.

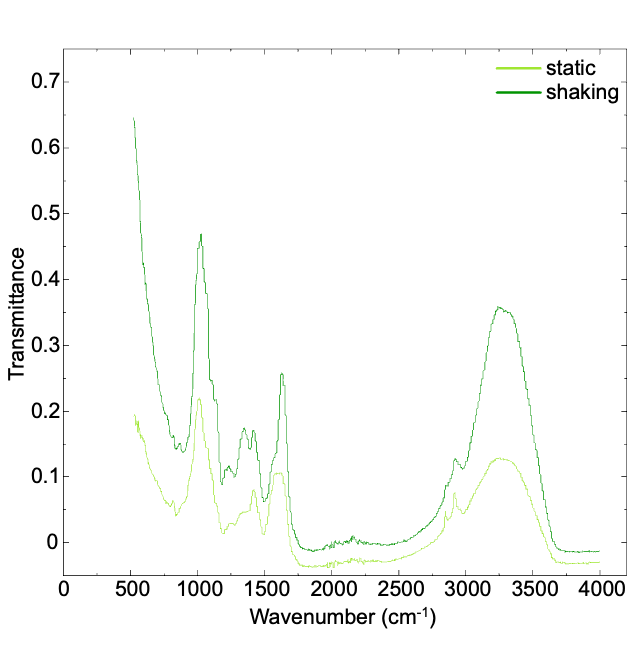

**Figure S4. Functional groups present in mycelia fiber with varied growth conditions.** Fourier Transform Infrared Spectroscopy spectra of a mycelia fiber grown using either static (light green) or shaking (dark green) growth.

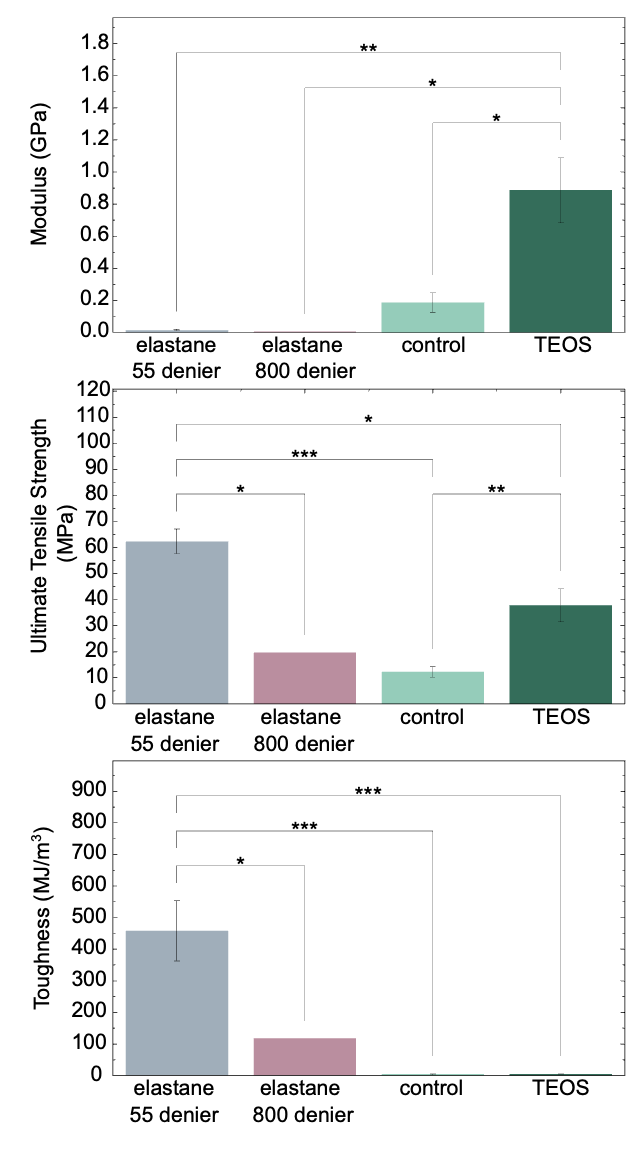

**Figure S5. Comparison of mycelia fiber to synthetic textile fiber.** Modulus, ultimate tensile strength, and toughness of fibers tested by uniaxial tensile testing. Elastane 55 denier (n=3, grey) and elastane 800 denier (n=1, pink) tested as received. Mycelia fiber control (n=5, light green) and mycelia TEOS (n=4, dark green). The data represent the mean and standard error of the mean. * *P* < 0.05, ** *P* < 0.01, *** *P* < 0.001, one-way ANOVA followed by Tukey’s *post hoc* test.
